# Histone acetyltransferase activity of CREB-binding protein is essential for synaptic plasticity in *Lymnaea*

**DOI:** 10.1101/2021.05.26.445902

**Authors:** Dai Hatakeyama, Hiroshi Sunada, Yuki Totani, Takayuki Watanabe, Ildikó Felletár, Adam Fitchett, Murat Eravci, Aikaterini Anagnostopoulou, Ryosuke Miki, Takashi Kuzuhara, Ildikó Kemenes, Etsuro Ito, György Kemenes

## Abstract

In eukaryotes, CREB-binding protein (CBP), a coactivator of CREB, functions both as a platform for recruiting other components of the transcriptional machinery and as a histone acetyltransferase (HAT) that alters chromatin structure. We previously showed that the transcriptional activity of cAMP-responsive element binding protein (CREB) plays a crucial role in neuronal plasticity in the pond snail *Lymnaea stagnalis*. However, there is no information on the role CBP plays in CREB-initiated plastic changes in *Lymnaea*. In this study, we characterized the *Lymnaea* CBP (LymCBP) gene and investigated the roles it plays in synaptic plasticity involved in regulating feeding behaviors. Similar to CBPs of other species, LymCBP possesses functional domains, such as KIX domain, which is essential for interaction with CREB and was shown to regulate long-term memory (LTM). *In situ* hybridization showed that the staining patterns of LymCBP mRNA in the central nervous system were very similar to those of *Lymnaea* CREB1 (LymCREB1). A particularly strong LymCBP mRNA signal was observed in the Cerebral Giant Cell (CGC), an identified extrinsic modulatory interneuron of the feeding circuit, key to both appetitive and aversive LTM for taste. Biochemical experiments using the recombinant protein of LymCBP HAT domain showed that its enzymatic activity was blocked by classical HAT inhibitors such as curcumin, anacardic acid and garcinol. Preincubation of *Lymnaea* CNSs with these HAT inhibitors blocked cAMP-induced long-term potentiation between the CGC and the follower B1 motoneuron. We therefore suggest that HAT activity of LymCBP in the CGCs is a key factor in synaptic plasticity contributing to LTM after classical conditioning.

## Introduction

*De novo* gene transcription is required for consolidation of long-term memory (LTM) (Kandel, 2001) and cAMP-responsive element binding protein (CREB)-dependent gene expression is one of the crucial steps of this process in mammals (Matos et al., 2019; Laviv et al., 2020) as well as invertebrates (Lakhina et al., 2015; Zhou et al., 2015; Hirano et al., 2016). Transcriptional activation requires the recruitment of multifunctional coactivators, resulting in a variety of different types of epigenetic modifications of histones and genomic DNA (Cedar and Bergman, 2009). In CREB-dependent control of gene expression, CREB-binding protein (CBP) functions as a coactivator of CREB. To date, CBP has been reported to be critical for LTM consolidation in mammals (Alarcón et al., 2004; Korzus et al., 2004; Chatterjee et al., 2013) and the gastropod *Aplysia* (Guan et al., 2002; Zhou et al., 2015). One of the most important functions of CBP is due to its histone acetyltransferase (HAT) activity, which stimulates gene transcription (Martinez-Balbás et al., 1998). Several studies have shown that histone acetylation by CBP is necessary for hippocampal long-term potentiation (LTP) (Korzus et al., 2004; Vecsey et al., 2007). Pharmacological inhibition of CBP HAT activity using the inhibitors curcumin and garcinol was reported to block memory consolidation (Zhao et al., 2012; Monsey et al., 2015; Merschbaecher et al., 2016) and memory-associated neuronal plasticity (Maddox et al., 2013a). These studies demonstrated that HAT activity of CBP plays an essential role in regulating synaptic plasticity associated with memory consolidation.

The pond snail *Lymnaea stagnalis* is a widely used organism to understand evolutionarily conserved molecular mechanisms of the consolidation of LTM for taste-related associations (Kemenes et al., 2006; Hatakeyama et al., 2013a; Murakami et al., 2013; Totani et al., 2020; Nakai et al., 2020a; 2020b). Sadamoto et al. (2004) first succeeded in the cloning of an isoform of CREB from the central nervous system (CNS) of *Lymnaea* and defined it as LymCREB1. They identified 7 different isoforms of LymCREB1 by alternative splicing and found that aversive taste conditioning significantly increased LymCREB1 gene expression (Sadamoto et al., 2010). Ribeiro et al. (2003) showed that appetitive taste conditioning selectively increased phosphorylated LymCREB1 in the ‘learning ganglia’ (the cerebral and buccal ganglia) of the *Lymnaea* CNS.

An identified extrinsic modulatory neuron type of the feeding system, the cerebral giant cell (CGC) was reported to play crucial roles in LTM after both aversive and appetitive taste conditioning (Kemenes et al., 2006; Ito et al., 2012; Nikitin et al., 2013). Injection of LymCREB1 siRNA into CGCs reduced the amplitude of excitatory postsynaptic potential (EPSP) in monosynaptic follower neurons of the CGCs, the B1 motoneurons (Wagatsuma et al., 2006), suggesting that regulation of gene expression by LymCREB1 was required for synaptic enhancement in memory consolidation. Based on these previous findings, we hypothesized that HAT activity of the *Lymnaea* CBP (LymCBP) plays an important role in the LymCREB1 initiated molecular processes of synaptic consolidation.

In the present study, we first cloned the LymCBP cDNA from the *Lymnaea* CNSs and identified neurons expressing the LymCBP mRNA. Focusing on the CGC, we pharmacologically and electrophysiologically analyzed the relationship between the HAT activity of LymCBP and the synaptic plasticity involved in the aversively conditioned feeding behavior of *Lymnaea*. Our findings provide new insights into the functions of LymCBP in synaptic plasticity underlying the consolidation of associative LTM.

## Materials and Methods

### Molecular cloning of LymCBP

To clone LymCBP, a series of degenerate PCR was performed with TaKaRa Ex Taq^®^ (Takara Clontech) and primers, which were designed at the basis of highly conserved domains, such as Taz1 (transcriptional <u>a</u>dapter <u>z</u>inc-binding 1), KIX domain and Taz2 domain, of *Aplysia* CBP (ApCBP; GenBank accession number: AY064470). After the sequential analyses of the Taz1, KIX and Taz2 domains of LymCBP, we performed 5’ and 3’RACE, and the PCR for the internal region between Taz1 and KIX domains and between KIX and Taz2 domains. The FirstChoice® RLM-RACE Kit (ThermoFisher) was used for amplification of 5’ and 3’ ends of LymCBP. All PCR products were subcloned with TA Cloning^®^ Kit (ThermoFisher) or pGEM^®^-T Easy Vector System (Promega). Nucleotide sequences of primers were summarized in Table 1 (No. 1-14).

**Table 1.**
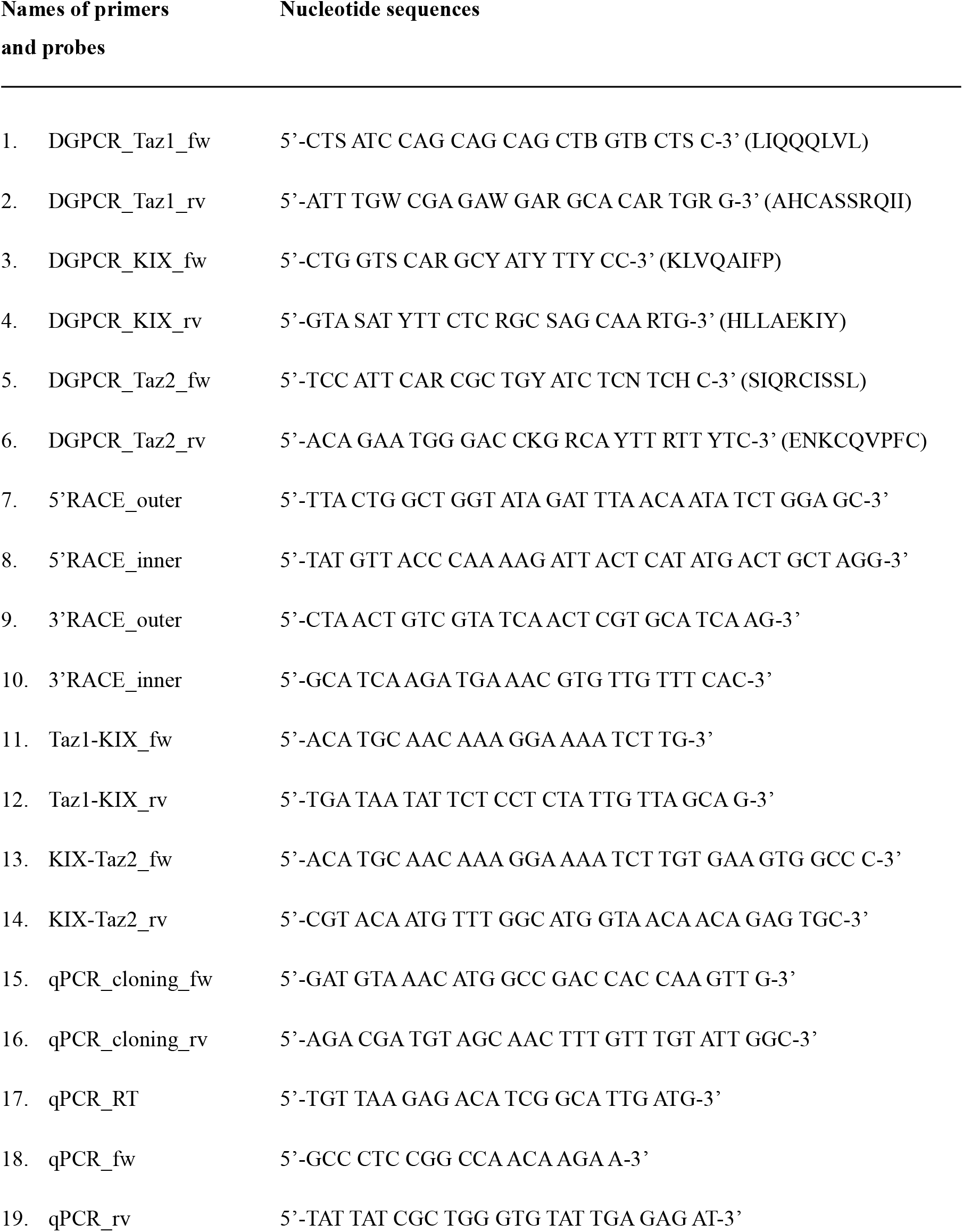
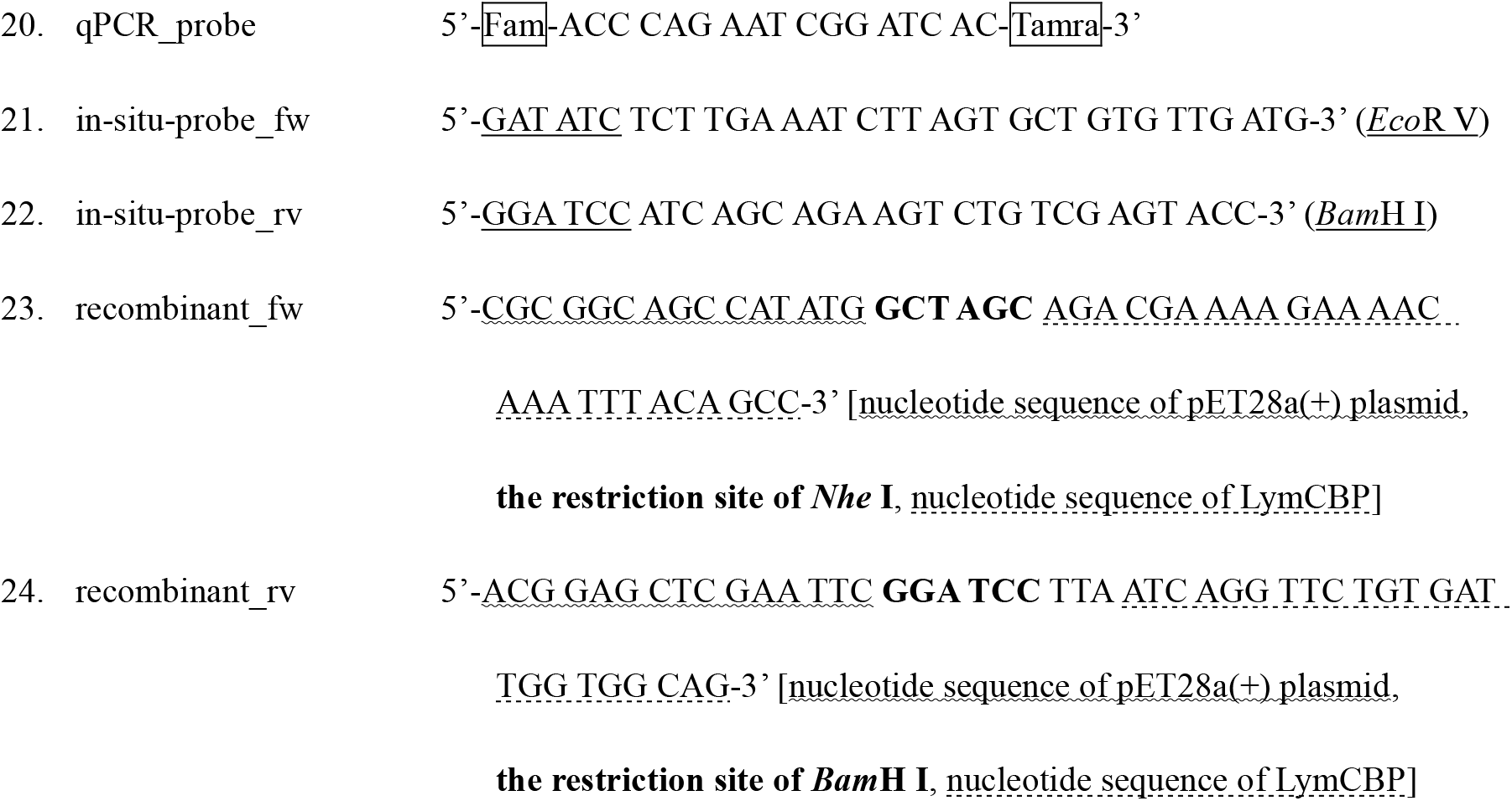
Nucleotide sequences of primers used for cloning (No. 1-14), synthesis for *in situ* hybridization probe (No. 15-16), synthesis of recombinant protein (No. 17-18), and qRT-PCR (No. 19-24) of LymCBP. Amino acid sequences of ApCBP for designing degenerate PCR (DGPCR) was shown with nucleotide sequences.

### Phylogenetic tree of LymCBP

To construct a phylogenetic tree of the CBP/p300 proteins, we aligned the full-length deduced amino acid sequence of the *Lymnaea* CBP with those of the known homologs of other species (listed with each Accession Number in Table 2) by using the MUSCLE algorisms (Edgar, 2004) on the Geneious (v9.1) program (available from http://www.geneious.com/). Maximum likelihood tree was constructed from the aligned sequences using the MEGA 6 program (Tamura et al., 2013) with default settings of the program. 1000 bootstrap replications were conducted to evaluate the reliabilities of the reconstructed trees. The obtained tree was visualized with the FigTree (v1.4.2) program (available from http://tree.bio.ed.ac.uk/software/figtree/). A CBP homolog of the Choanoflagellate *Salpingoeca rosetta* was used as an outgroup.

**Table 2.**
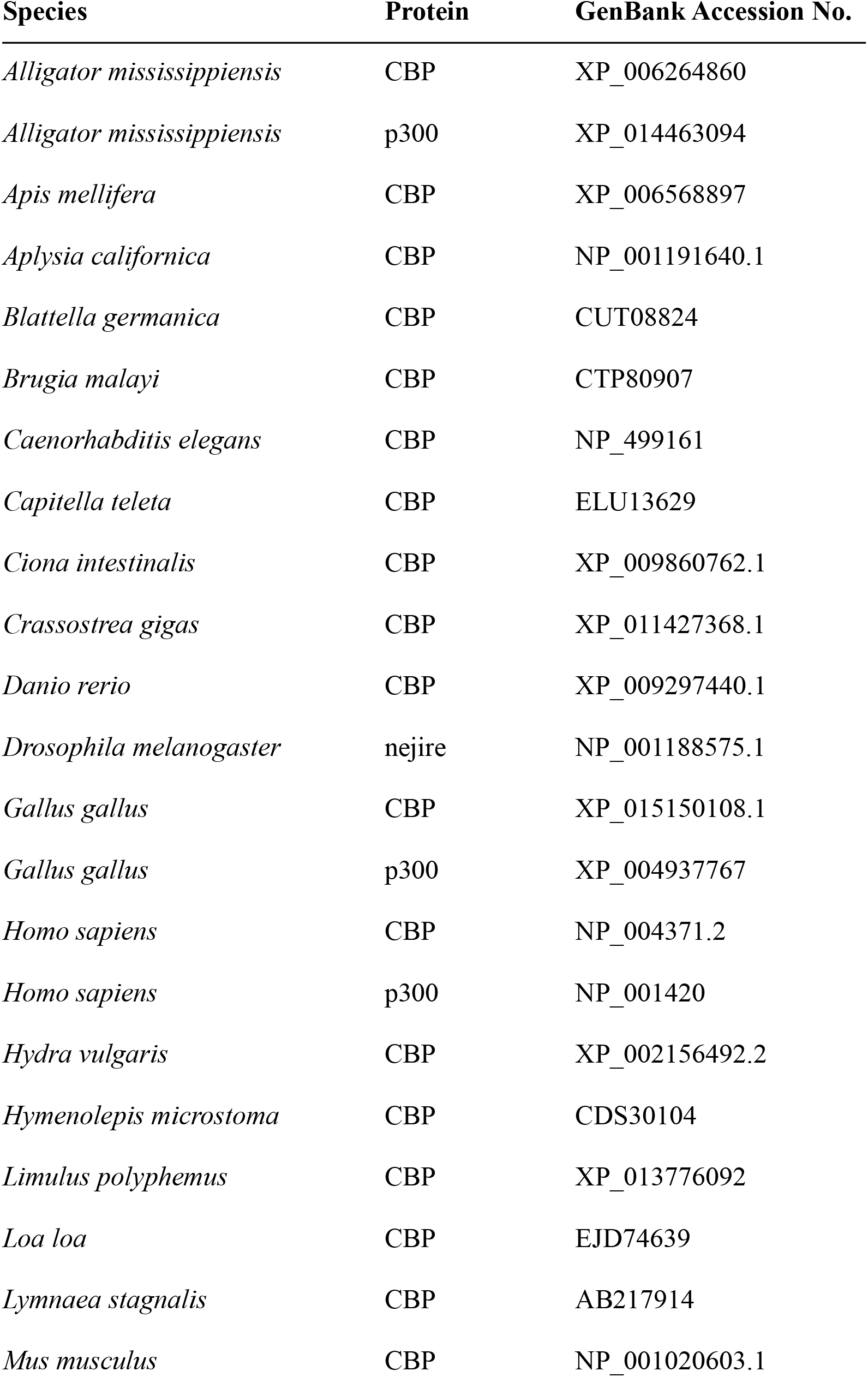
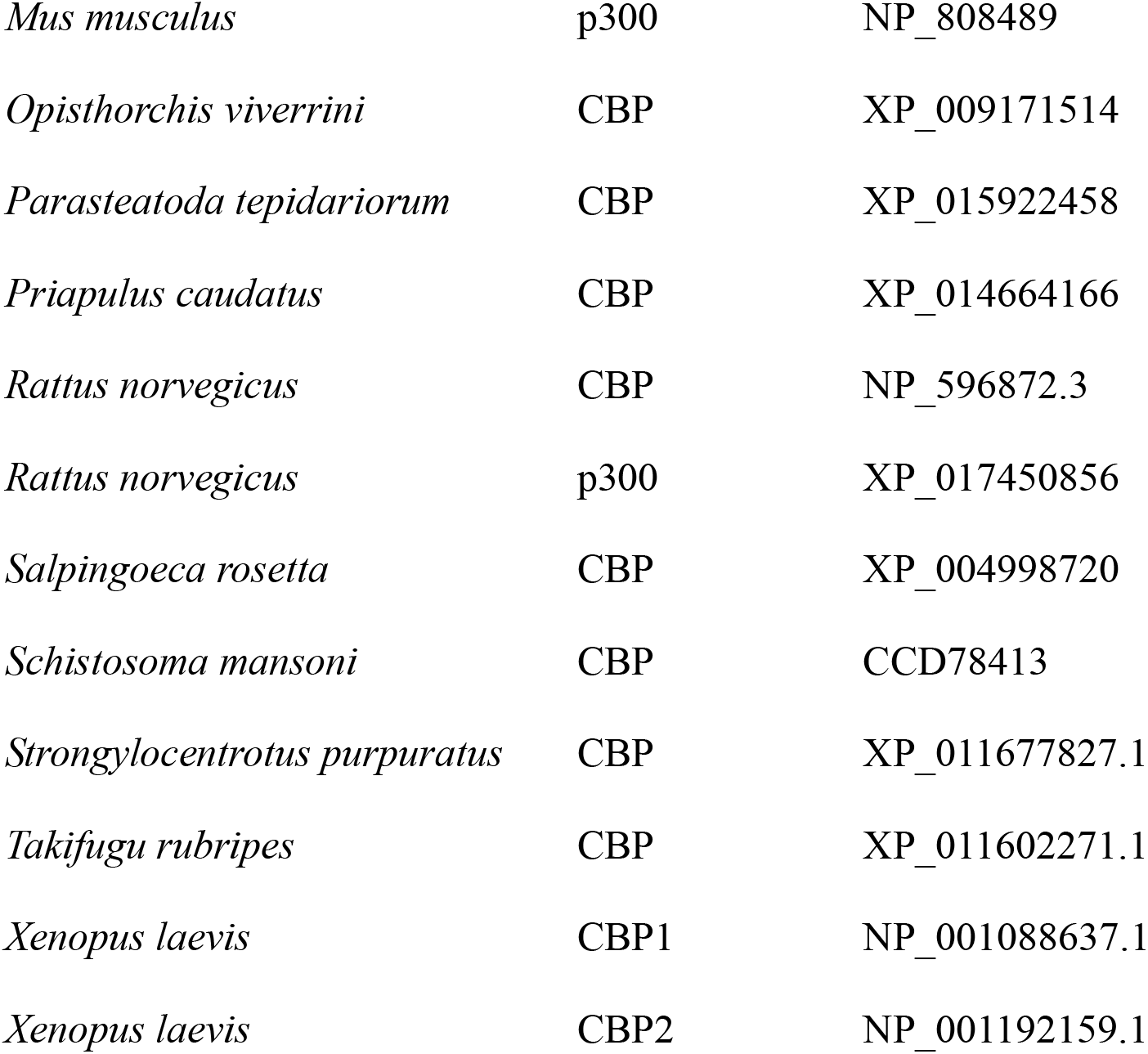
GenBank accession numbers of various CBP and p300 used for molecular phylogenetic tree shown in Fig. 2.

### TaqMan-based quantitative PCR (qPCR)

To quantify the absolute copy number of LymCBP mRNA in several *Lymnaea* tissues, we performed TaqMan-based qPCR by establishing RNA standard curves derived from a dilution series of known template concentrations. The procedures for this experiment were modified from previous reports (Hatakeyama et al., 2004b, 2006; Wagatsuma et al., 2005). Partial LymCBP gene (from 303rd to 722nd nucleotides, which contains the open reading flame of the *N*-terminal region) was amplified by RT-PCR with primers shown in Table 1 (No. 15, 16), and the PCR product was subcloned into pGEM^®^-T Easy Vector (Promega). The plasmid was purified with Plasmid Midi Kit (QIAGEN) and digested with restriction enzyme *Spe* I. The LymCBP RNA was synthesized with MAXIscript^TM^ (ThermoFisher), purified with RNeasy^®^ Mini Kit (QIAGEN), quantified the concentration, and used as the standard RNA. Serially diluted standard RNA (5 × 10^1^ – 5 × 10^6^ copies/μl) were reverse-transcribed in 10 μl of reaction mixture to prepare the first strand cDNA with 2.5 μM of the specific primer for LymCBP diluted to 1:5 and used as the standard cDNAs.

Total RNA from different types of *Lymnaea* tissue (CNS, buccal mass, penis, ovotestis, gut and mantle) was purified with TRIzol^TM^ Reagent (ThermoFisher). Reverse transcription (RT) of standard series (5×10^1^ – 5×10^6^ copies) and total RNA of *Lymnaea* tissues (each 300 ng) were performed in the buffer containing 0.2 μg yeast tRNA, 1× PCR Buffer II (ThermoFisher), 5.5 mM MgCl_2_, 0.5 mM dNTP, 0.2 μM gene specific primer for LymCBP, 0.3 unit/ml Prime RNase inhibitor (Eppendorf), and 0.9 units/ml MultiScribe^TM^ Reverse Transcriptase (ThermoFisher) for 10 min at 25°C, for 30 min at 48°C, and for 5 min at 95°C. All RT samples were diluted to 5 times by adding RNase-free distilled water. The diluted RT solution of samples and standards (1 μl) were added to PCR-reaction solution [final concentration: 1× TaqMan^®^ Gene Expression Master Mix (ThermoFisher), 50 nM each primer, and 50 nM probe]. Nucleotide sequences of primers and probes were summarized in Table 1 (No. 17-20). Reaction was carried out at 95°C for 10 min, 45 cycles of 95°C for 15 sec, and 60°C for 1 min with real-time PCR system (StepOnePlus Real-Time PCR System, ThermoFisher). In the assay, several doses of standard cDNA (1×10^1^ – 1×10^6^ copies) were applied in triplicate to estimate inter-assay coefficients of variation between runs.

In situ *hybridization of LymCBP.* The procedures for this experiment were modified from previous reports (Hatakeyama et al., 2004a; 2006; 2010). Non-functional region (from 278th to 1146th of nucleotide sequence) of LymCBP was isolated by RT-PCR (see Table 1 (No. 21, 22) for nucleotide sequences of primers). After insertion of this PCR product into pGEM^®^-T Easy Vector, the plasmid was linearized with *Eco*R V and *Bam*H I to generate digoxigenin-labeled antisense and sense RNA probes, respectively. The procedures for generating the probes, hybridization and coloring were the same as those that were previously reported (Hatakeyama et al., 2004a; 2006).

### Western blotting, co-precipitation and mass spectrometry

#### Experimental animals

Pond snails*, Lymnaea stagnalis*, were bred at the University of Sussex, Brighton, United Kingdom and were maintained in large holding tanks containing copper-free water at 18-20^0^C on a 12-hour light - dark cycle. The animals were fed on a vegetable-based fish food (Tetra-Phyll; TETRA Werke, Melle, Germany) twice a week and lettuce three times a week.

#### Preparation of snail CNS samples for Western blotting, co-precipitation and mass spectrometry

The whole CNS of four-month-old snails were dissected. Briefly, the shell of the snail was removed, and the preparation was then pinned down in a Sylgard-coated dish containing HEPES buffered saline and dissected under a stereomicroscope (E-Zoom6, Edmund Optics, Barrington NJ, USA). The CNS was accessed by a dorsal body incision and isolated from the buccal mass by severing all the peripheral nerves and immediately placed in Eppendorf tubes on dry ice and stored at −80°C.

#### Western blotting

*Lymnaea* CNS lysates were prepared in RIPA buffer. Protein (10-20 µg) was electrophoresed on homemade 6% Bis-Tris protein gels and transferred to polyvinylidene difluoride (PVDF) membrane. The polyclonal antibody to anti-human-CBP (C20, sc-583, Santa Cruz) was used at 1:500 dilution in 4% skimmed milk in TBS-Tween overnight at 4°C with rotation. Following washes with TBS-Tween (10 min ×3), the membranes were incubated with the anti-rabbit-IgG conjugated with horseradish peroxidase (Sigma Aldrich) at room temperature at 1:10,000 dilution for 2 hr. Finally, the membranes were cleaned from the excess of secondary antibody by washing with TBS-Tween (10 min ×3) and TBS (10 min ×1). Detection of blot was carried out with a LumiSensor^TM^ HRP Substrate Kit (GenScript Technology).

#### Immunoprecipitation assay and immunoblotting

Sixty pooled *Lymnae*a CNSs were homogenised in RIPA lysis buffer supplemented with 2 mM PMSF, 1 mmol/L Na_3_VO_4_ and protease inhibitors, (Santa Cruz, sc-24948) by using a micropestelle. 50 μl of the pre-immunoprecipitated lysate was saved as “input” to be used for immunoblotting. The homogenate was pre-cleared in a Pierce Spin column (Thermofisher Scientific, 69705) containing 40 μl of Protein A/G PLUS-Agarose (Santa Cruz, sc-2003) for one hour at 4^0^C on a rotor. Four Pierce Spin columns containing 40 μl of Protein A/G PLUS-Agarose suspension were prepared. Three of the Pierce Spin columns were used to prepare Antibody/Agarose complex and one of the columns was used as a negative control without containing the antibody. 500 μl of HEPES buffer was added to all of the Protein A/G PLUS-Agarose spin columns. For the Antibody/Agarose complex preparation, 10 μg of anti-mouse CREB2 antibody (Santa Cruz, sc-390063) was added to each of the Protein A/G PLUS-Agarose Spin column. The columns were incubated on a rotor at RT for 3 hrs. The columns were centrifuged at 2500 RPM for 1 min and the flow-through was removed. The columns were then washed/centrifuged four times at 2500 RPM with HEPES buffer at 4^0^C. After the final wash, 500 μg of CNS homogenised lysate was transferred into each column containing the Antibody/Agarose complex and the one column without containing the antibody. The columns were placed on a rotor and incubated overnight at 4^0^C. The following day, the columns were centrifuged at 2500 RPM for 1 min and the lysate was removed. The columns were washed/centrifuged four times at 2500 RPM with HEPES buffer. The eluates for each column were eluted in 50 μl of Glycine, pH 2.8. The pH of the eluates was neutralized with 5 μl of 1M Tris, pH 9.5. The IP products were pooled together and 100 μl of the IP product was used for LC/MS analysis and 45 μl of the IP product was prepared for Western Blotting. 45 μl of the IP product was resuspended in 15 μl of 4 x Loading sample buffer and boiled at 90^0^C for 10 mins. Immunoprecipitates, negative control and 25 μg of the pre-immunoprecipitated lysate (5% input) were resolved in 10% SDS-polyacrylamide gel and transferred to a nitrocellulose membrane. Non-specific binding sites were blocked with 5% milk in TBS solution with 0.1% Tween-20 (1xTBST) for 1 hour at RT. Membranes were incubated with anti-mouse CREB2 antibody (1:1000, Santa Cruz, sc-390063) overnight at 4^0^C. The following day, the membrane was washed three times for 10 mins in TBST and incubated with highly cross-adsorbed Infra-Red Dye 680RD goat anti-mouse secondary antibody (1:5000, LI-COR, 926-68070) for 1 h at RT, followed by two washes with 1xTBST and a final wash of 1xPBS. Membranes were developed using the LI-COR Odyssey Fc Fluorescence Imaging system.

#### Protein precipitation for detergent removal

In order to remove the detergents used in our immunoprecipitation protocol, which are incompatible with the downstream MS analysis we have used the methanol/chloroform precipitation method as previously described (Wessel and Flugge, 1984). Protein precipitates were then resuspended in 50 μl of 5 X Invitrosol^TM^ LC/MS Protein Solubilizer. The samples were incubated at 60°C for 5 mins, vortexed and incubated at 60°C for another 10 mins. 200 μl of 25 mM ammonium bicarbonate (ABC Buffer) was added per sample in order to obtain an Invitrosol^TM^ concentration of 1X. The samples were separated into 5 tubes containing 50 μl of sample per tube.

#### In-solution Trypsin digestion with the MS detergents

Samples were reduced in 10 mM Dithiothreitol (DTT) for 30 mins at RT and cysteines were alkylated in the dark in 60 mM Iodoacetamide (IAA) for 20 min at RT. 2 μl of Trypsin/Lys-C mix (0.5μg/μl, Promega, V5073) was added to each sample and incubated for 3 hrs at RT in the dark. After 3 hrs, 150 μl of ABC buffer per tube was added and incubated overnight at RT. The following day, the digestion was stopped by adding 200 μl of 5% Acetonitrile, 3% Trifluoroacetic acid. Tryptic peptides were desalted with stage tips as previously described (Rappsilber et al., 2003) The desalted peptides were concentrated in a Speed-Vac for 10 mins at low power. The peptides were reconstituted with 15 μl of 5% Acetonitrile, 0.1% Formic Acid and transferred to a microplate for LC/MS analysis.

#### LC/MS analysis

Peptides were separated by reverse-phase chromatography using a Dionex Ultimate 3000 nanoLC (Dionex/Thermo Fisher Scientific) on a C18 capillary column (Acclaim PepMap100 C18, 2 μm, 100 Å, 75 μm i.d.×20 cm / Thermo Fisher Scientific) at an eluent flow rate of 300 nl/min. Mobile phase A contained 5% acetonitrile, 0.1% formic acid and mobile phase B contained 80% acetonitrile 0.1% formic acid. The column was preequilibrated with 5% mobile phase B followed by an increase to 65% mobile phase B in 75 min. The eluted peptides were ionized online by electrospray ionization (ESI) and transferred into an LTQ Orbitrap XL mass spectrometer (Thermo Fisher Scientific) which was operated in the positive mode to measure full scan MS spectra (300–1700 m/z in the Orbitrap analyzer at resolution R = 30,000) followed by isolation and fragmentation (MS/MS) of the ten most intense ions (in the LTQ iontrap) by collision-induced dissociation (CID).

#### MS/MS data analysis

The analysis of the raw MS and MS/MS Spectra was performed using the MaxQuant software (version. 1.6.3.4). Initial maximum precursor and fragment mass deviations were set to 7 ppm and 0.5 Da, respectively. Variable modification (oxidation of methionine and N-terminal acetylation) and fixed modification (cysteine carbamidomethylation) were set for the search against a UniProt FASTA formatted database of the taxonomic clade *euthyneura* (including *Lymnaea stagnalis*). Trypsin with a maximum of two missed cleavages was chosen for digestion. The minimum peptide length was set to 7 amino acids and the false discovery rate (FDR) for peptide and protein identification was set to 0.01.

### Recombinant protein of LymCBP

The partial recombinant protein of the LymCBP HAT domain (from 1240th arginine to 1641st lysine, GenBank accession number BAE02656.1) with His_6_-tag was synthesized in *Escherichia coli* and purified as previously described with slight modifications (Hatakeyama et al., 2014, 2018a). LymCBP catalytic domain was amplified by RT-PCR with cDNA prepared from *Lymnaea* CNSs, PrimeSTAR^®^ HS DNA Polymerase (Takara Clontech) and primers, whose nucleotide sequences were summarized in Table 1 (No. 23, 24). The PCR product was ligated into pET28a(+) plasmid (Novagen), which was linearized with *Nhe* I and *Bam*H I, using In-Fusion^®^ HD Cloning Plus kit (Takara Clontech). Then, the Rosetta-gami^TM^ 2(DE3) competent cells (Novagen) was transformed with this ligated plasmid. Expression of the partial recombinant protein of the LymCBP catalytic domain was induced by treatment with 0.32 mM of isopropyl β-D-thiogalactopyranoside (IPTG) in TBG-M9 medium [0.8% Bacto Tryptone (BD) and 0.4% NaCl], followed by purification using nickel-nitrilotriacetic acid (Ni-NTA) agarose resin (QIAGEN). The recombinant protein was desalted with PD-10 column (Sephadex^TM^ G-25M, GE Healthcare), purified using a cation exchange column [HiTrap carboxymethyl (CM) FF column (GE Healthcare)] installed in the ÄKTAprime plus system (GE Healthcare), and condensed with Amicon^®^ Ultra-15 column (Merck Millipore).

### Acetylation assays using radioisotope-labeled acetyl-CoA

The procedures followed to perform this experiment were modified from previous reports (Hatakeyama et al., 2018a; 2018b). The recombinant LymCBP HAT domain (200 ng) were incubated with the histone H1 purified from calf thymus (Sigma-Aldrich, 1 μg) and 7.4 kBq of [^14^C]-acetyl-CoA (Moravek, Inc.) for 4 hr in buffer containing 50 mM Tris-HCl (pH 8.0), 10% glycerol, 1 mM dithiothreitol, and 10 mM sodium butyrate. Curcumin (MedChemExpress), anacardic acid (MedChemExpress) and garcinol (Enzo) prepared in dimethyl sulfoxide (DMSO) were premixed at each 50 μM. Reactions were separated in 14% SDS-PAGE gels. To detect β radiation released from radiocarbon, the gels were dried on pieces of filter papers and exposed on imaging plates for several days. Signals were observed using a fluoroimage analyzer (FLA-2000; Fuji Film).

### Intracellular recording

The whole CNS, including the cerebral ganglia and the buccal ganglia was dissected out from the snail in *Lymnaea* saline [50 mM NaCl, 1.6 mM KCl, 2.0 mM MgCl_2_, 3.5 mM CaCl_2_, and 10 mM HEPES (pH 7.9)]. The isolated CNS was immobilized on a Sylgard-coated dish using stainless-steel pins. In order to physically separate the cerebral giant cells and B1 motoneurons, a Vaseline dam was constructed around the cerebral ganglia. This enabled us to apply a solution that contains the inhibitors to the CGCs. The CGC and the B1 motoneuron were impaled with glass microelectrodes filled with 3 M KCl giving tip resistances of 20–50 MΩ. A train of 5 spikes in the CGC, which was produced by a current injection (0.8–1.8 nA) for 1 sec, evoked a single large compound EPSP in the B1 motoneuron (electric stimulator: SEN-7203, Nihon Kohden, Tokyo, Japan; intracellular recording amplifier: MEZ-8300, Nihon Kohden; AD converter: DIGIDATA 1322A, Axon Instruments, Foster City, CA, USA). We used the neurons located in the ipsilateral side.

### Injection of cAMP or 5’AMP to the CGC

Based on a previous study (Nakamura et al., 1999), cyclic AMP (cAMP) or 5’AMP (Sigma-Aldrich), which was filled into a glass microelectrode (50–100 MΩ) as a 200 mM solution dissolved in 20 mM Tris buffer (pH7.5), was injected into the CGC by passing hyperpolarizing current pulses (50 msec on, 50 msec off) of 4 nA for 20 min. Before and after the injection of hyperpolarizing current into the CGC, the EPSP recorded in the B1 motoneuron is not changed by activation of the CGC. The data were recorded in the same preparation at 30 min and 3 hr after cAMP or 5’AMP injection. The data at 0 hr (i.e., pre) were recorded before cAMP injection. For the analysis of EPSP changes recorded in the B1 motoneuron, the peak amplitude of the EPSP was measured.

### Application of HAT inhibitors to the CGC

We used the following three different inhibitors of HATs: curcumin (Sigma), anacardic acid (Sigma) and garcinol (Enzo). Each of these drugs was dissolved in DMSO as a stock solution at 67.9 mM (curcumin), 50 mM (anacardic acid) and 20 mM (garcinol). Stock solutions were diluted 1000-fold in *Lymnaea* saline then applied to the CGC area where the cerebral ganglia were surrounded by Vaseline wall and incubation for 1 hr. After incubation with each drug, intracellular recordings for the measurement of EPSPs were made. The data was expressed as the mean ± SEM. Significant differences at *p* < 0.05 between pre-injection, 30 min and 3 hr after injection cAMP or 5’AMP were examined by one-way repeated measures ANOVA and post hoc Tukey’s test. For the normalized EPSP amplitude, the values after 30 min and 3 hr for each group compared to the pre-injection were examined with one-way repeated measured ANOVA and post hoc Tukey’s test.

## Results

### Properties *of the* LymCBP gene and protein

The arrangement of the functional domains in LymCBP was the same as that of other p300/CBPs (Fig. 1A). The full sequence of LymCBP (GenBank accession number: AB217914) was about 75% identical to *Aplysia californica* CBP (ApCBP; GenBank accession number: AAL54859) in amino acid sequence level. In functional domains, such as Taz1 domain, KIX domain, bromodomain, HAT domain, and Taz2 domain of LymCBP, the amino acid sequences were individually over 90% identical to those of ApCBP and human CBP (GenBank accession number: U47741) (Fig. 1B-D). Comparing amino acid sequences among LymCBP, human CBP and human p300, the functional amino acid residues in bromodomain for interacting with acetylated histones (Dhalluin et al., 1999; Mujtaba et al., 2004; Plotnikov et al., 2014; Xu et al., 2017), those in HAT domain for intramolecular interactions (Delvecchio et al., 2013; Ortega et al., 2018; Zhang et al., 2018), and interaction with acetyl-CoA (Liu et al., 2008; Maksimoska et al., 2014) were highly conserved (Table S1). Molecular phylogenetic analysis of the p300/CBP proteins revealed that *Lymnaea* CBP is closely related to CBP proteins found in other molluscan species such as *Aplysia* and the oyster *Crassostrea giga* (Fig. 2).

**Fig. 1.**
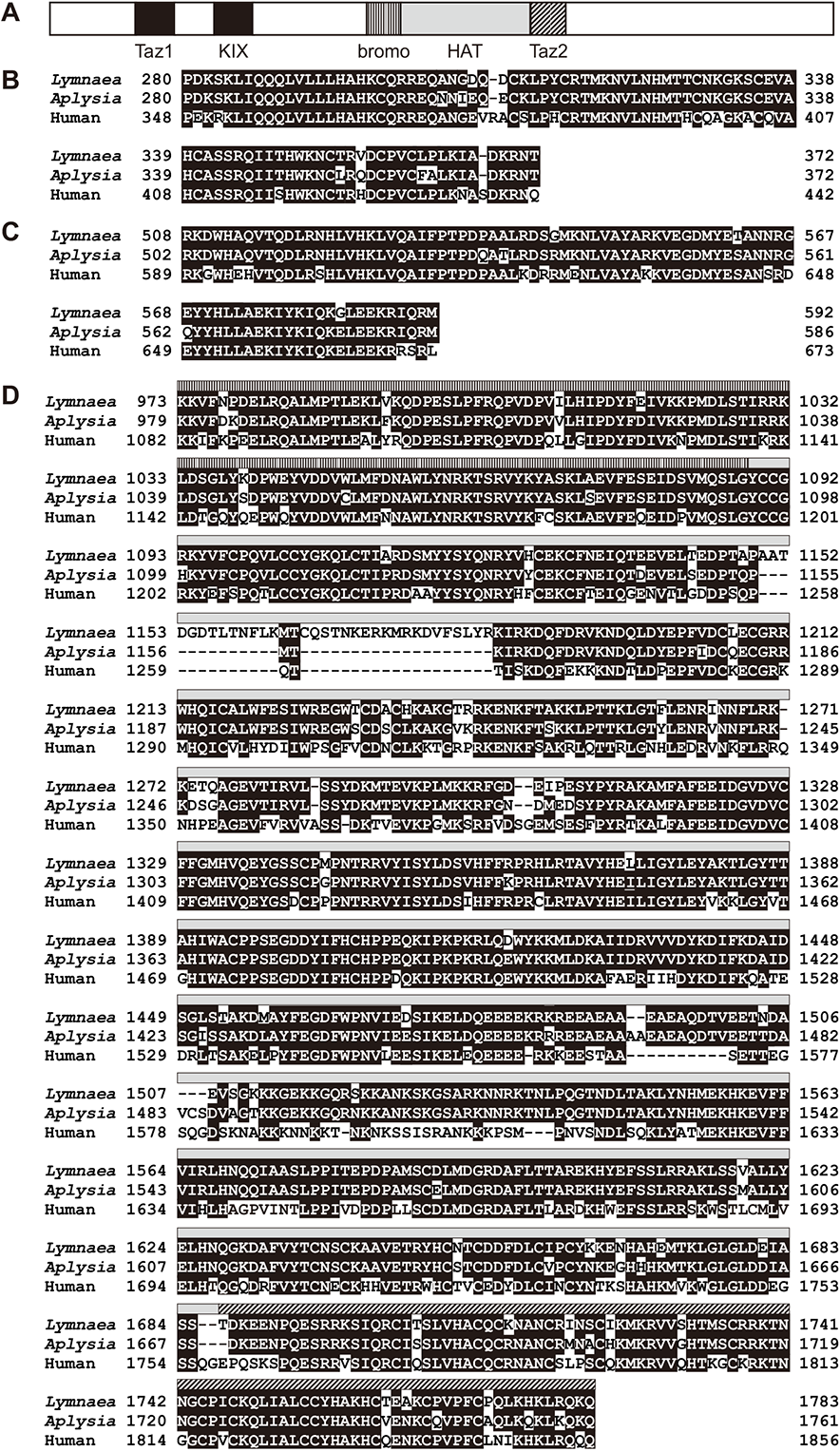
Predicted amino acid sequence of *Lymnaea* CBP. (A) Diagram of the structure of *Lymnaea* CBP (GenBank accession number: AB217914). Main functional domains (Taz1 domain, KIX domain, bromodomain, acetylation domain and Taz2 domain) are highlighted with patterned bars. Alignments of amino acid sequences of (B) Taz1 domain, (C) KIX domain and (D) compound of the bromodomain, HAT domain and Taz2 domain between *Lymnaea* CBP and *Aplysia* CBP (GenBank accession number: AY064470) and human CBP (GenBank accession number: U47741). The patterns of bars in (D) correspond to those shown in (A).

**Fig. 2.**
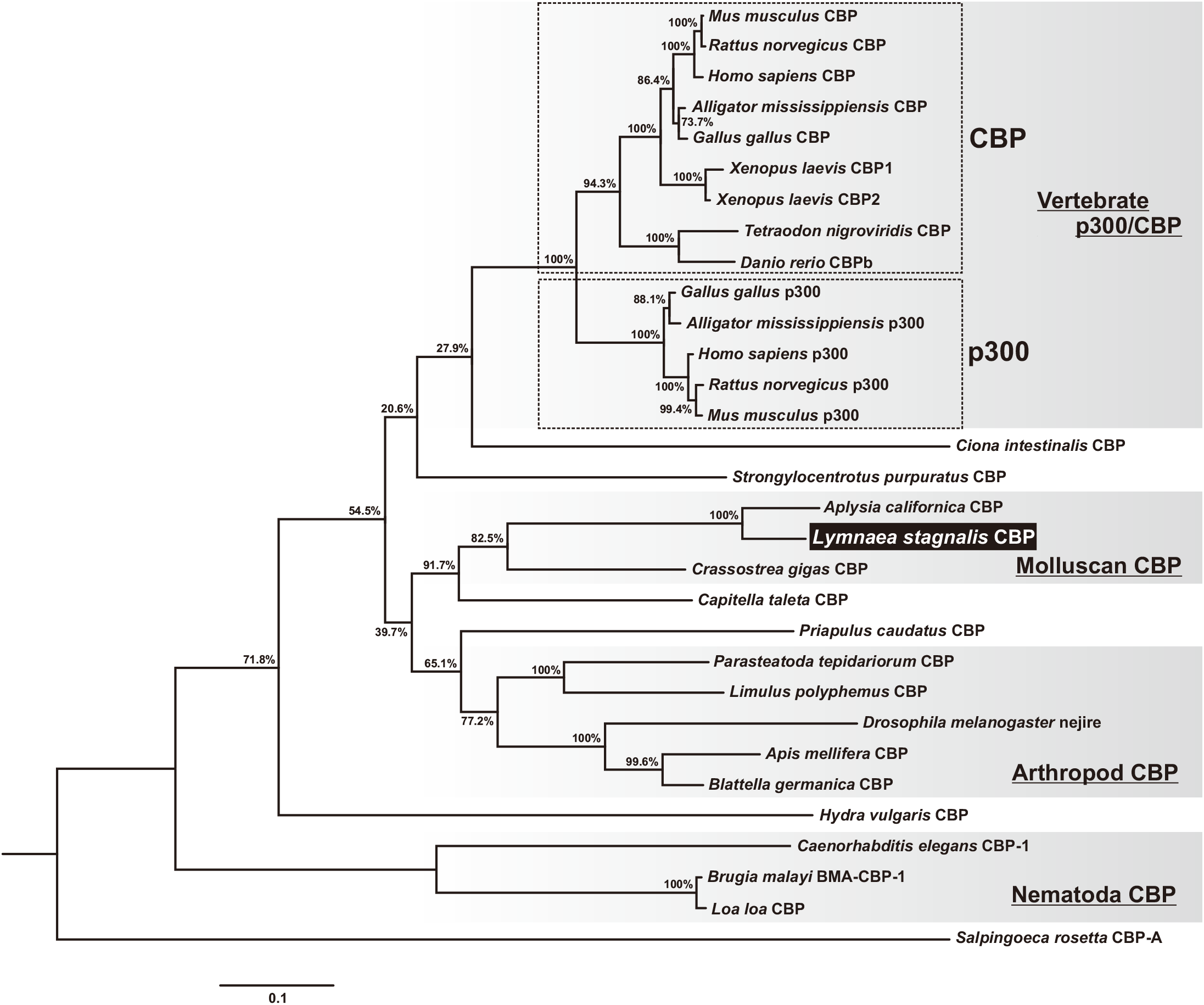
Molecular phylogenetic tree of *Lymnaea* CBP and other p300/CBP proteins. The scale bar indicates 0.1 substitutions per site. Bootstrap values are shown at the nodes. A CBP homologue of the Choanoflagellate *Salpingoeca rosetta* was used as an outgroup. The GenBank accession numbers of p300/CBP-like proteins are listed in Table 1.

### Localization of LymCBP mRNA in the *Lymnaea* CNS

Absolute copy numbers of LymCBP mRNA in whole CNS and other tissues were analyzed by quantitative PCR (qPCR) (Fig. 3). More than 15,000 copies of LymCBP mRNA were contained in 6 ng of total RNA of CNSs (Fig. 3). This value was 1.4 to 2.9 times higher compared to other tissues (Fig. 3).

**Fig. 3.**
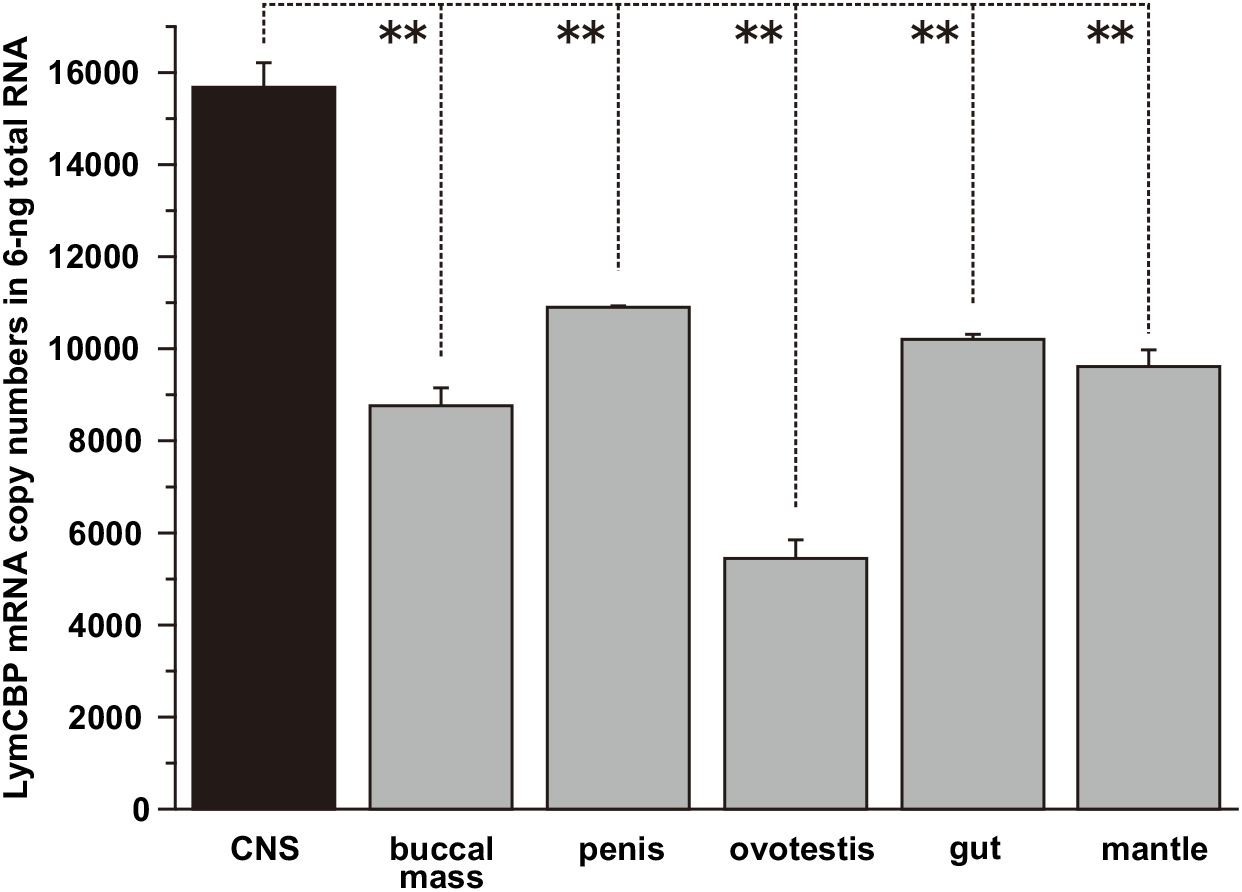
Quantification of LymCBP mRNA in the whole CNS of *Lymnaea*. LymCBP is more abundantly transcribed in the CNS than other tissues of *Lymnaea*. The data are expressed as the mean ± SEM; *p* < 0.01, \*\**p* < 0.01, Tukey’s post-hoc test.

*In situ* hybridization showed the localization of LymCBP mRNA in the *Lymnaea* CNSs (*n* = 8). Weak intensity of the positive signal was detected in some neurons in the buccal ganglia (Fig. 4A). The neurons localized at the ventral edge of the ganglia seemed to be parts of the B4 cluster (B4CL) cells. Strong signal was observed in the CGCs in the cerebral ganglia (Fig. 4B,C). The same signal was also seen in identifiable large neurons, the RPeD1 (Fig. 4D) and the LPeD1 (Fig. 4E), in the pedal ganglia. Notably, the staining patterns of LymCBP mRNA in the buccal, cerebral and pedal ganglia were very similar to those of LymCREB1 (Sadamoto et al., 2004). The same positive signal of LymCBP was also detected in unidentified neurons in parietal and visceral ganglia (Fig. 4F). No positive signals were detected in negative control experiments with sense LymCBP cRNA probe (data not shown).

**Fig. 4.**
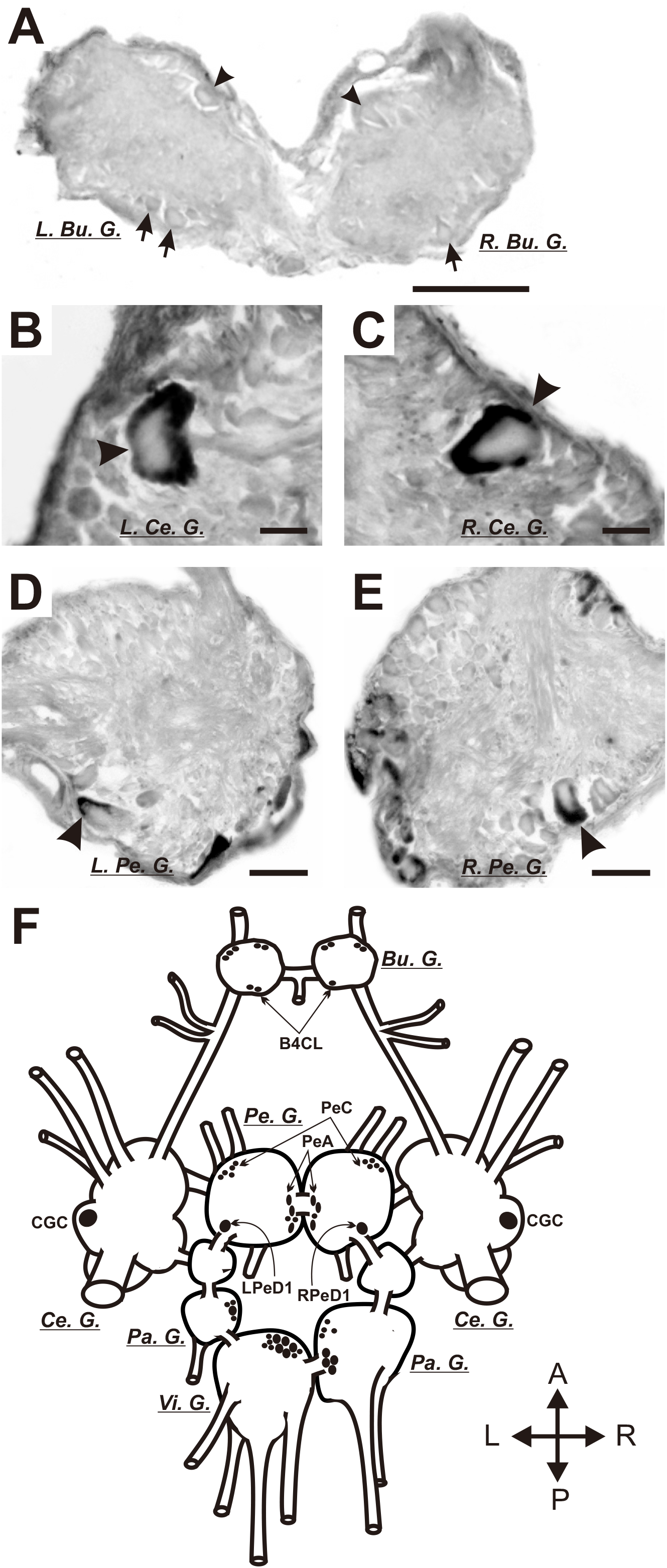
Localization of LymCBP mRNA in identifiable neurons shown by *in situ* hybridization. (A) Weak signal of LymCBP could be observed in several neurons (arrowheads), especially in the B4CL feeding motoneurons (arrows), in the buccal ganglia. Bu. G.: buccal ganglion, L: left; R: right. (B-E) Strong signal of LymCBP mRNA was observed in both CGCs (arrowheads in B and C), RPeD1 (arrowhead in D) and LPeD1 (arrowhead in E). Ce. G.: cerebral ganglion, Pe. G.: pedal ganglion. (F) Schematic drawing of the distribution of neurons containing LymCBP mRNA. Pa. G.: parietal ganglion, Vi. G.: visceral ganglion. Scale bars = 100 μm (A, D and E) and 20 μm (B and C).

### Existence of the protein form of LymCBP

Western blotting demonstrated the existence of the protein form of LymCBP in the CNS. The epitope of the anti-human-CBP antibody is the *C*-terminal region of human CBP, which is highly conserved in LymCBP (Fig. 1D). The western blotting method detected two positive bands (Fig. 5A). The larger band, indicated with a filled arrowhead around the marker of 245 kDa, was considered to be LymCBP, whose molecular weight was estimated as 252.3 kDa from the amino acid sequence (Fig. 1). The smaller band shown with an open arrowhead may be an isoform of LymCBP (Fig. 5A). In the EMBL-EBI database, twenty transcript variants of CBP are found in human, and, of these isoforms, seven transcripts can be translated (Gene: “CREBBP”, ENSG00000005339). Therefore, it is suggested that *Lymnaea* also expresses the smaller isoform of CBP.

**Fig. 5.**
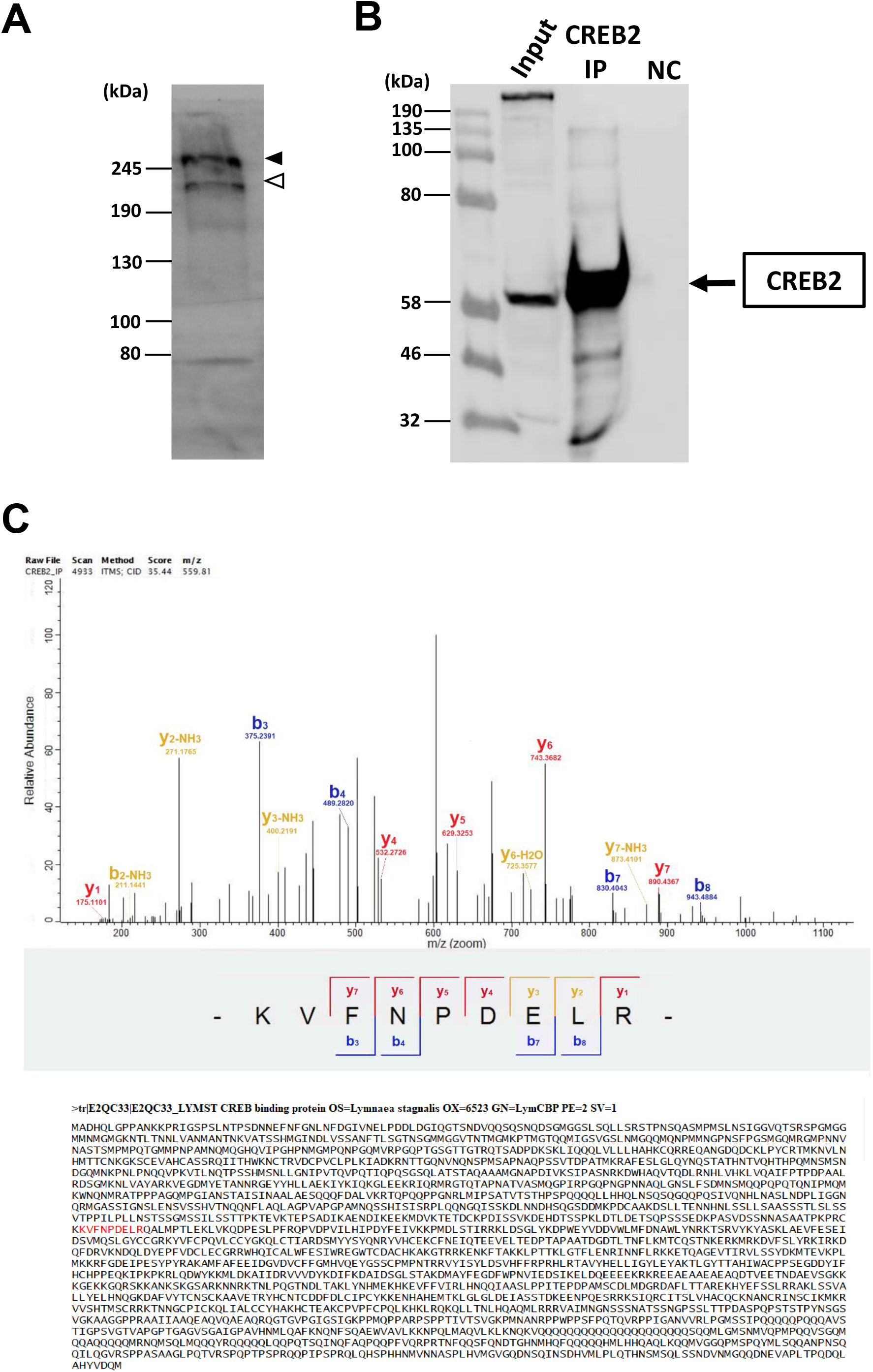
Identification of the protein form of LymCBP. (A) Existence of LymCBP protein in CNSs was confirmed by western blotting. Molecular weight of the positive band (245 kDa; filled arrowhead) was consistent with theoretical value calculated from amino acid sequence of LymCBP (252.3 kDa). A putative smaller isoform of LymCBP was also detected (open arrowhead). (B, C) CREB-Binding Protein identified as an interacting partner of CREB2 by MS/MS analysis in the CNS of *Lymnaea stagnalis*. (B) Sixty extracted CNS of *Lymnaea stagnalis* were pooled, homogenised and subjected to immunoprecipitation using a specific CREB2 antibody. Immune complexes were immunoblotted for CREB2. (C) MS/MS Spectrum of the CBP identified as one of the interacting partners of CREB2 in the CNS of *Lymnaea stagnalis*. CREB2 IP product was subjected for MS/MS analysis and a CBP specific peptide for *Lymnaea* CBP (KVFNPDELR) was identified.

Phosphorylation of CREB1 at Ser 133 is triggering its kinase inducible domain (KID) to bind to the KID interacting domain (KIX) of CREB binding protein (CBP) (Mayr and Montminy, 2001; Radhakrishnan et al., 1997; Shaywitz et al., 2000). To identify the interaction of CBP with CREB proteins in our model organism *Lymnaea stagnalis* we have utilized immunoprecipitation (IP) and liquid chromatography mass spectrometry (LC/MS).

Although in pilot experiments we found that none of our CREB1 antibodies worked well for IP assays with total CNS lysates from *Lymnaea stagnalis*, previous studies have shown that human CREB2 (human activating transcription 4, hATF4), a transcriptional repressor of CREB1, can directly interact with multiple domains of the CBP including i. the KIX domain, ii. a region that contains the third zinc finger and the E1A-interacting domain, iii. the C-terminal region containing the p160/SRC-1-interacting domain and iii. the histone acetyltransferase domain which can directly interact with histone acetyltransferase p300 transcriptional coactivator of CBP (Liang and Hai, 1997). In addition, CBP and p300 can acetylate CREB2 in its bZIP domain and enhance its transcriptional activity (Gachon et al., 2002; Lassot et al., 2005; Liang and Hai, 1997). In order to confirm these reported interactions of CBP and CREB2, we have performed IP assays with a specific antibody for CREB2 and CNS homogenates from *Lymnaea stagnalis* (Fig. 5B), followed by identification of the CREB2 interaction partners by LC/MS (Fig. 5C). As shown in Fig. 5C, after LC/MS, the data analysis of the CID fragmentation spectrum from a peptide with the mass of 559.81 m/z revealed the amino-acid sequence of a unique peptide (KVFNPDELR) from the UniprotKB entry E2QC33 of *Lymnaea stagnalis*, validating that *Lymnaea* CBP is an interacting partner of CREB2 in the CNS. Importantly, these results further confirmed that LymCBP is expressed in the *Lymnaea* CNS.

### Effects of HAT inhibitors on LymCBP enzymatic activity

To investigate the enzymatic properties of LymCBP as a functional HAT, we produced a partial recombinant protein of its HAT domain (from 1240th to 1641st, Fig. 1D) and investigated its acetyltransferase activity. The recombinant protein, to which an His_6_-tag was attached at its *N*-terminus, could be purified by precipitation using Ni-NTA agarose resin. To remove contaminated proteins derived from bacterial cells, the recombinant protein solution was desalted and then purified by cation-exchange chromatography. This purification procedure was successful, and only the bands of the recombinant protein were observed in the 12th to 19th fractions (arrowhead in Fig. 6A). However, the concentrations of these preparations were too low to be measured by Bradford protein assay. Then, the eight fractions were mixed and treated with a centrifugal filter to condense the recombinant protein, resulting its concentration increased to around 80 ng/μL (lane 2 in Fig. 6B), which was sufficient for the biochemical experiments.

**Fig. 6.**
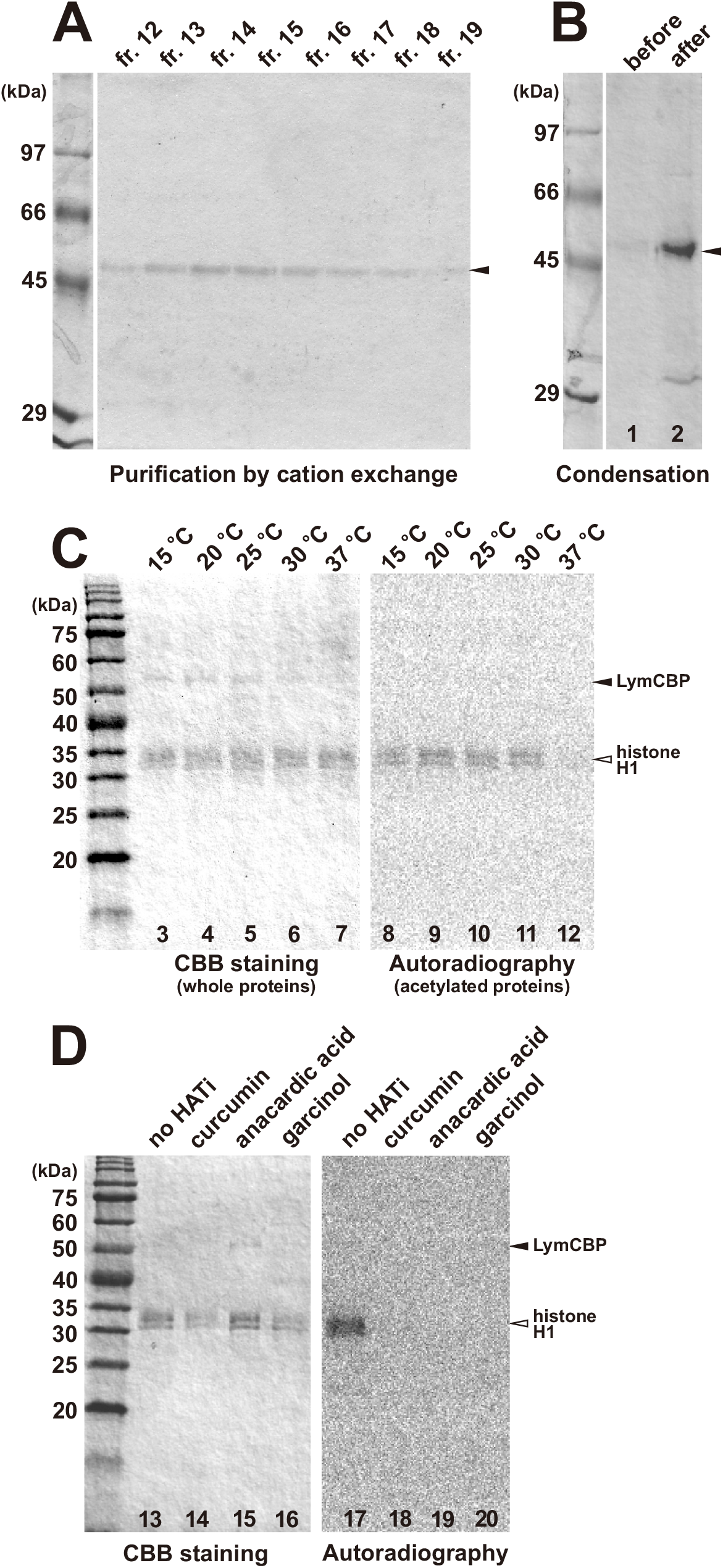
HAT activity of LymCBP was blocked by inhibitors. (A) The partial recombinant protein of LymCBP HAT domain purified by cation-exchange chromatography was detected in fractions No. 12–19 (bands shown with arrowhead). The concentrations of these preparations were too low to be measured by Bradford protein assay. (B) All fractions obtained by cation-exchange chromatography were mixed (lane 1), and the recombinant protein was condensed using a centrifugal filter. A high enough concentration of the recombinant protein for biochemical experiments was obtained (lane 2). (C) Condensed recombinant LymCBP HAT domain was incubated with histone H1 at different temperatures; 15°C, 20°C, 25°C, 30°C and 37°C. CBB staining of the gel showed that same amount of histone H1 was loaded (open arrowhead, lanes 3-7). However, the band of recombinant LymCBP HAT domain (filled arrowhead) was drastically decreased after incubation at 37°C, suggesting that LymCBP was denatured (lane 7). A positive signal of acetylation was observed after incubation at 15°C, 20°C, 25°C and 30°C (lanes 8-11). This signal was almost eliminated after incubation at 37°C (lane 12). (D) Effects of the HAT inhibitors curcumin, anacardic acid and garcinol. Reaction mixtures were incubated at 30°C. Although the same amount of recombinant LymCBP HAT domain (filled arrowhead) and histone H1 (open arrowhead) was incubated, CBB staining showed that the band signals of histone H1 were attenuated after incubating with curcumin (lane 14) and garcinol (lane 16), suggesting that these inhibitors degraded the protein. Autoradiography indicated that, without any HAT inhibitor, positive signal of acetylation was detected on the band of histone H1 (lane 17). This signal was eliminated by addition of each HAT inhibitor (lanes 18-20), showing that the acetylation activity of LymCBP recombinant protein was significantly blocked by HAT inhibitors.

We first determined the incubation temperature for the *in vitro* acetylation assay. A reaction mixture containing recombinant LymCBP HAT domain, bovine histone H1, and [^14^C]-acetyl-CoA were incubated for 4 hr at different temperatures; 15°C, 20°C, 25°C, 30°C and 37°C. Separation by SDS-PAGE and Coomassie Brilliant Blue (CBB) staining of the gel showed that same amount of histone H1 was loaded (lanes 3-7 in Fig. 6C). However, the band of recombinant LymCBP HAT domain was notably decreased after incubation at 37°C, suggesting that LymCBP was degraded (lane 7 in Fig. 6C). By autoradiography, the positive signal indicating acetylation of histone H1 was observed when incubated at 15°C, 20°C, 25°C and 30°C (lanes 8-11 in Fig. 6C), and this signal was almost eliminated after incubation at 37°C (lane 12 in Fig. 6C). Therefore, we decided to incubate samples at 30°C in following acetylation assays.

Next, the effects of HAT inhibitors, namely curcumin, anacardic acid and garcinol, on the HAT activity of LymCBP was investigated. After adding each HAT inhibitor (each at 50 μM final concentration) or DMSO as negative control, samples were incubated for 4 h at 30°C and separated by SDS-PAGE. Although the amount of each protein in each tube should be same, CBB staining showed that the band intensities of histone H1 were decreased by incubating with curcumin and garcinol (lanes 14 and 16 in Fig. 6D), suggesting that these inhibitors degraded the protein. Autoradiography showed that LymCBP acetylated histone H1 without any HAT inhibitors (lane 17 in Fig. 6D). In the presence of HAT inhibitors, LymCBP HAT activity was significantly blocked and acetylation signals were eliminated (lanes 18-20 in Fig. 6D). These results showed that similar to human CBP, LymCBP had HAT activity, and this enzymatic activity was significantly blocked by HAT inhibitors.

### Synaptic plasticity between a buccal motoneuron and its regulatory interneuron induced by cAMP

The isolated *Lymnaea* CNS was incubated with DMSO saline for 1 h before the injection of cAMP into the CGC cell body. We tested the changes in the EPSP in the B1 motoneuron by activation of the CGC after the cAMP injection. We measured the amplitudes of the compound EPSPs 30 min and 3 h after the onset of cAMP injection. The CGC was depolarized by current injection to fire action potentials, evoking the compound EPSP in the B1 motoneuron. We used a a train of 5 spikes in the CGC to evoke large compound EPSPs in the B1 motoneuron because the single EPSPs in the B1 motoneuron were very small (approximately 1 mV). These compound EPSPs at 30 min and 3 h after in preparations with cAMP-injected CGCs were about twice as large as pre injection EPSPs [Fig. 7A, *F*(2,18) = 10.35, pre injection vs 30 min: *p* < 0.05; pre injection vs 3 hr: *p* < 0.05, *n* = 10]. In contrast, there were no significant changes in the 5’AMP-injected preparations [Fig. 7B, *F*(2,12) = 1.192, pre injection vs 30 min: *p* > 0.05; pre injection vs 3 hr: *p* > 0.05, *n* = 7]. Thus, the effect of HAT inhibitors on the compound EPSPs 30 min and 3 hr after cAMP injection were examined in the following experiments.

**Fig. 7.**
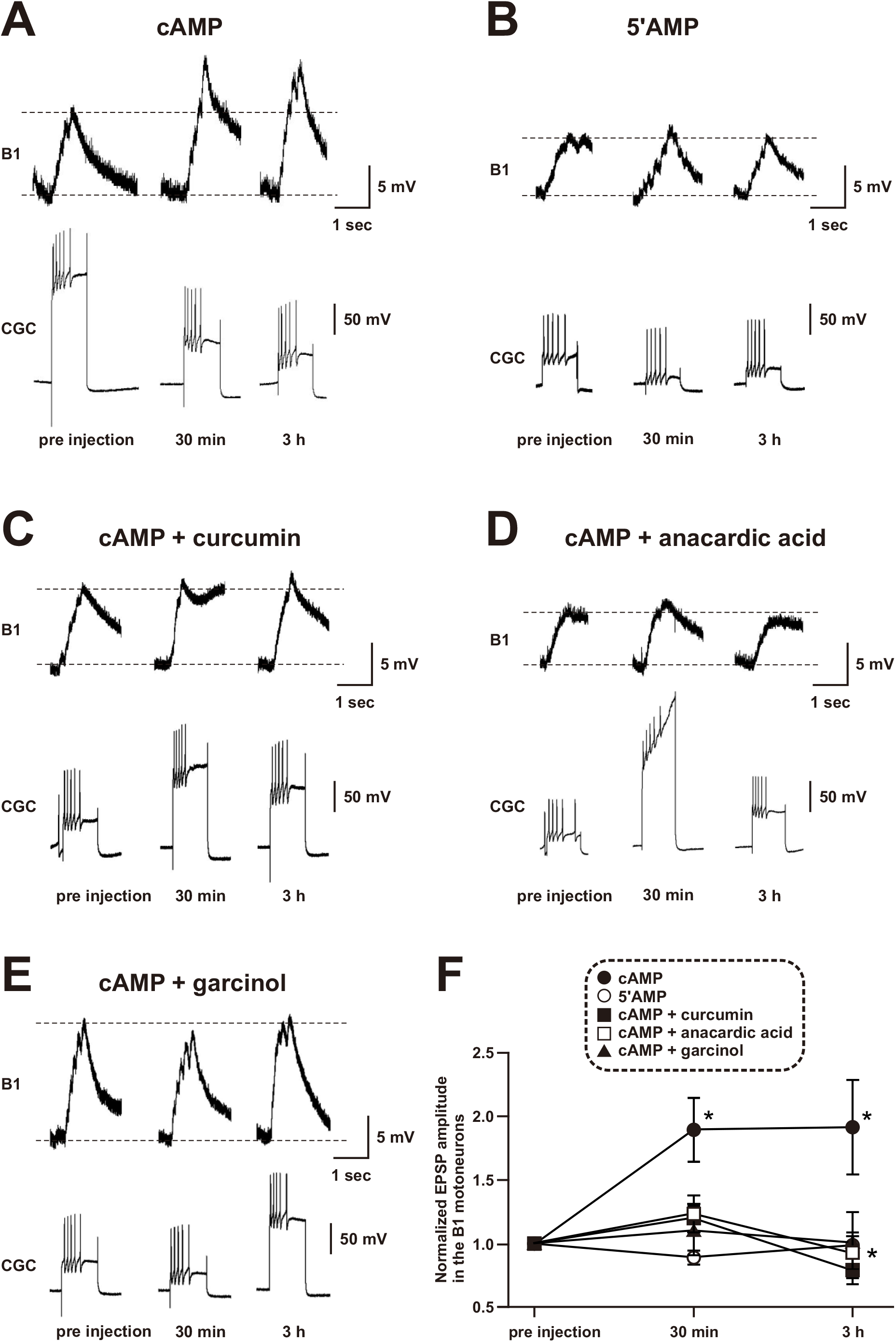
Long-term synaptic changes at synapses between the CGC and the B1 motoneuron in the isolated CNS are blocked by HAT inhibitors. EPSPs were evoked in the B1 motoneuron by the injection of depolarizing current into the CGC. The isolated CNS was incubated with DMSO saline or HAT inhibitors for 1 h prior to injection of cAMP. (A) cAMP injection into the CGC cell body increased EPSP amplitude in B1 motoneuron at both 30 min and 3 h after the injection. (B) In contrast, EPSP amplitude increase was not observed in the group which was injected with 5’AMP. (C, D, E) The CNS was incubated with saline containing one of the HAT inhibitors curcumin (67.9 μM), or anacardic acid (50 μM), or garcinol (20 μM) for 1 h prior to injection of the cAMP. In these groups, the EPSP was not increased either at 30 min or 3h after the injection of cAMP. (F) The enhancement of B1 EPSPs is shown as the summarized data. Injection of cAMP into the CGC cell body increased EPSP amplitude in the B1 motoneuron, but pre-treatment with either HAT inhibitor did not elicit enlargement of EPSP. The data are expressed as the mean ± SEM, \**p* < 0.05, Tukey’s post-hoc test.

### Effects of HAT inhibitors on the EPSP in the B1 motoneuron

HAT inhibitors were shown to possess significant inhibitory effects on HAT activity of LymCBP (Fig. 6C). Therefore, we examined the effects of the application of these HAT inhibitors to CGCs on the compound B1 EPSPs that were found to be increased after cAMP injection into the CGC. As reported previously (Nakamura et al., 1999; Sadamoto et al., 2004), the EPSP in the B1 motoneuron is facilitated by injection of cAMP into the CGC (Fig. 7A). This increase in EPSP amplitude was not observed when 5’AMP was injected into CGCs instead of cAMP (Fig. 7B). Next, we investigated changes to the cAMP-induced EPSP in the presence of the HAT inhibitors curcumin, anacardic acid and garcinol. Incubation with 67.9 µM curcumin of the cerebral ganglion for 1 h did not change the amplitude of the compound EPSPs in the B1 motoneuron (Fig. 7C). After 1-hr incubation with curcumin prior to the cAMP injection there was no significant change in the amplitude of the compound EPSPs 30 min and 3 hr after the cAMP injection compared with pre injection [Fig. 7C, *F*(2,22) = 6.001, pre injection vs 30 min: *p* > 0.05; pre injection vs 3 hr: *p* > 0.05, *n* = 12]. However, a significant difference was observed between 30 min and 3 hr. The incubation with 50 µM anacardic acid on the cerebral ganglion for 1 hr that did not enlarge of the compound EPSPs in the B1 motoneuron (Fig. 7D). The 1-hr incubation with anacardic acid prior to the cAMP injection, in this group, there was no significant differences in the amplitude of the compound EPSPs 30 min and 3 hr after the cAMP injection compared with pre injection [Fig. 7D, *F*(2,10) = 1.563, pre injection vs 30 min: *p* > 0.05; pre injection vs 3 hr: *p* > 0.05, *n* = 6]. The incubation with 20 µM garcinol on the cerebral ganglion for 1 hr that did not enlarge of the compound EPSPs in the B1 motoneuron (Fig. 7E). The incubation for 1 hr with garcinol prior to the cAMP injection, in this group, there was no significant differences in the amplitude of the compound EPSPs 30 min and 3 hr after the cAMP injection compared with pre injection [Fig. 5E, *F*(2,14) = 0.01214, pre injection vs 30 min: *p* > 0.05; pre injection vs 3 hr: *p* > 0.05, *n* = 8].

The results from all these groups are summarized in Fig. 7F. The effect of HAT inhibitors on increasing compound EPSPs amplitude was tested by taking a 0 min baseline recording followed by a 30 min and a 3 hr recording after cAMP was injected. The compound EPSPs were normalized to the initial baseline value. Incubation with the HAT inhibitors before the injection of cAMP, did not result in increased compound EPSPs amplitudes, as seen in the DMSO saline treatment group. In the host hoc Tukey’s test, there were significant differences in the following comparisons: DMSO saline treatment cAMP injection group; pre injection vs 30 min; pre injection vs 3 hr, curcumin treatment group; 30 min vs 3 hr).

## Discussion

In the present study we, for the first time, identified a conserved CREB-binding protein in the CNS of *Lymnaea* (LymCBP) and showed the importance of its HAT activity for synaptic plasticity between an identified interneuron (CGC) and a postsynaptic motoneuron (B1) of the feeding system. The CGC is a key interneuron for consolidating both aversive and appetitive LTM for taste, and both LymCBP and LymCREB1 were transcribed in this neuron (Fig. 4). A previous report showed that inhibiting functions of LymCREB1 by pre-injection of excess amount of a CRE oligonucleotide and siRNA of LymCREB1 into the CGC blocked cAMP-induced synaptic facilitation between the CGCs and the B1 motoneurons (Sadamoto et al., 2004; Wagatsuma et al., 2006), and suggested a crucial role of LymCREB1 as a transcriptional factor in synaptic facilitation. Here, inhibition of LymCBP HAT activity using HAT inhibitors similarly blocked the synaptic potentiation (Fig. 7), suggesting that this synaptic facilitation requires the co-operation of LymCREB1 and LymCBP HAT activity.

### Gene structure of LymCBP

We succeeded in the cloning of the full-length cDNA of LymCBP from the *Lymnaea* CNS. Comparing its deduced amino acid sequence with those of *Aplysia* and human CBPs, several functional domains were found to be highly conserved (Fig. 1). This result suggests that LymCBP has similar functions to mammalian CBP and p300 (Fig. 2). The KIX domain of CBP associates with the kinase-inducible domain (KID) of CREB, and they construct a complex together (Chrivia et al., 1993). KID, which is also called as P-box, is highly conserved in LymCREB1 (Sadamoto et al., 2004). High conservation of the KIX domain in LymCBP and KID in LymCREB1 suggested that LymCBP and LymCREB could interact via these domains. Additionally, a previous report showed that the KIX domain of CBP is involved in the formation of spatial memory following hippocampus-dependent learning (Chatterjee et al., 2020). Here, we demonstrated that these two mRNAs were co-localized in the CGCs, key neurons for LTM consolidation (Fig. 4). Taken together, the consolidation of LTM in *Lymnaea* is suggested to be regulated by LymCBP and LymCREB1. The Taz1 and Taz2 domains of LymCBP were also highly conserved (Fig. 1). These domains function as interfaces to bind with a variety of transcription factors and HATs, such as Fos and p300/CBP-associated factor (pCAF) (Glass et al., 1997). It was previously reported that the bromodomain of mammalian CBP is important for acetylation of histones (Raisner et al., 2018) and interaction with acetylated histones (Zeng et al., 2008; Plotnikov et al., 2014). At least 15 amino acid residues in the bromodomain regulate interaction with acetylated histones (Dhalluin et al., 1999; Plotnikov et al., 2014; Xu et al., 2017), and, compared with amino acid sequences among human p300, human CBP and LymCBP, 14 out of 15 amino acid residues were conserved in LymCBP (Table S1). Other functional amino acid residues are highly conserved in the RING domain (Table S2), the acetyl-CoA-binding domain (Table S3), and the ZZ domain (Table S4). The RING domain in human p300/CBP has an autoinhibitory function for the HAT domain (Delvecchio et al., 2013; Ortega et al., 2019). The ZZ domain of human p300/CBP recognizes histone tails and modulates its enzymatic activity (Legge et al., 2004; Zhang et al., 2018). The most important domain in CBP is the KIX domain, which is well known as an interface for interacting with CREB (Kwok et al., 1994), and this domain is highly conserved in LymCBP (Fig. 1). These results suggested that not only the HAT activity, which was shown in Fig. 6, but also other functions of CBP are highly conserved in LymCBP.

Our immunoprecipitation assays indicated an interaction between LymCBP and LymCREB2 (Fig. 5B). A similar interaction also has been shown in the human homologues of these proteins (Liang and Hai, 1997). The interfaces in CBP for binding with CREB2 include the KIX domain, Taz2 domain and HAT domain (Liang and Hai, 1997). These domains were highly conserved in LymCBP (Fig. 1). In addition, acetylation of CREB2 by CBP was previously reported (Gachon et al., 2002). Therefore, the interaction between LymCBP and LymCREB2 shown in Fig. 5B may be needed for LymCREB2 acetylation by LymCBP.

### Expression of LymCBP in the Lymnaea CNSs

*In situ* hybridization showed that LymCBP mRNA was transcribed in a limited number of identifiable neurons (Fig. 4F). In particular, the CGC in the cerebral ganglia exhibited a strong LymCBP signal (Fig. 4B,C). Both mRNA and protein forms of LymCREB1 exist in the CGC (Ribeiro et al., 2003; Sadamoto et al., 2004), showing the co-localization of LymCBP and LymCREB1 in this neuron. The CGC was identified to be a key neuron for the long-lasting conditioned responses after taste aversion learning (Kojima et al., 2001; Yamanaka et al., 2000; Ito et al., 2012; Sunada et al., 2017) and appetitive classical conditioning (Kemenes et al., 2006; Nikitin et al., 2008; Vavoulis et al., 2010; Marra et al., 2013; Nikitin et al. 2013). The same signal was also seen in identified large neurons, the RPeD1 and the LPeD1, in the pedal ganglia (Fig. 4D,E). The RPeD1 was especially reported to be necessary for LTM formation in operant conditioning of aerial respiratory behavior (Scheibenstock et al., 2002). The staining patterns of LymCBP mRNA in the buccal, cerebral and pedal ganglia were very similar to those of LymCREB1 (Sadamoto et al., 2004), which possesses KID for interacting with LymCBP and plays a role in the enhancement of transcriptional activity (Sadamoto et al., 2004).

Our proteomics experiments confirmed the existence of translated LymCBP in the *Lymnaea* CNS, corroborating the findings from the DNA and RNA level assays of LymCBP expression.

In addition, we previously indicated the existence of both mRNA and protein forms of another transcriptional factor, CCAAT/enhancer binding protein of *Lymnaea* (LymC/EBP), in the CGC and the RPeD1 (Hatakeyama et al., 2004a). In mammalian cells, C/EBP is acetylated by p300/CBP, its acetylation boosts C/EBP-mediated transcriptional activity (Ceseña et al., 2007). Taken together, the interaction between LymCBP and transcriptional factors, such as LymCREB1 and LymC/EBP, may be involved in the consolidation of LTM of *Lymnaea*.

### Involvement of LymCBP HAT activity in synaptic plasticity

In this study, we used HAT inhibitors (curcumin, anacardic acid and garcinol; reviewed by Hatakeyama et al., 2013b) for biochemical and electrophysiological experiments (Figs. 6 and 7). Biochemical analyses using the recombinant LymCBP HAT domain showed that the HAT activity of LymCBP was significantly blocked by HAT inhibitors. Molecular docking simulation indicated the amino acid residues of the human p300, whose amino acid sequence of the HAT domain was highly identical with human CBP, for interaction with garcinol (Coste et al., 2020). For interaction between human p300 and garcinol, five amino acid residues (S1400, Y1414, D1444, Y1446 and Q1455) in its HAT domain were required, and these were completely conserved in LymCBP (S1356, Y1370, D1400, Y1402 and Q1411) (Table S5). Therefore, HAT activity of LymCBP might be inhibited by similar interaction mechanisms with garcinol.

Electrophysiological experiments showed that these chemicals inhibited cAMP-dependent synaptic plasticity between the CGCs and their follower neurons (Fig. 7). Although the block of synaptic plasticity by curcumin has not been reported in other species, its inhibitory effects on newly acquired or reactivated fear memories were indicated (Monsey et al., 2015). Similarly, garcinol was also reported to block newly acquired memories in mammals (Zhao et al., 2012; Maddox et al., 2013a; Monsey et al., 2017) and honeybees (Merschbaecher et al., 2016). Another HAT inhibitor, C646, which was not used in this study, significantly impaired memory consolidation (Maddox et al., 2013b; Merschbaecher et al., 2016). These inhibitory effects might be induced by blocking HAT activities of CBP and other HATs. On the other hand, there are many reports showing that curcumin has ameliorative effects from the impairment of memory and synaptic plasticity caused by amyloid peptides (Ahmed et al., 2011; Zhang et al., 2015), chronic stress (Liu et al., 2014), drugs (Soukhaklari et al., 2018), heavy metals (Namgyal et al., 2020), aging (Belviranli et al., 2013; Cheng et al., 2013) and viral proteins (Tang et al., 2009; Shen et al., 2015), suggesting that other molecular mechanisms, except for inhibiting HAT activity, rescue memory and synaptic plasticity.

## Acknowledgements

This work was partly supported by The Japan Society for the Promotion of Science (JSPS), a Grant-in-Aid for Scientific Research (C) to D.H. (No. 26460562 and 17K08867), Takeda Science Foundation to D.H, Waseda University Grants for Special Research Projects (2018K-141 and 2020C-135) to E.I. Work at the University of Sussex, UK, was funded by grants from the Medical Research Council to G.K. (MRC/G0400551), and the Biotechnology and Biological Sciences Research Council (BBSRC) to G.K. and I.K. (BB/H009906/1) and to I.K. and G.K. (BB/P00766X/1), respectively.

## Author Contributions

Experiments performed in the UK: D.H., I.F., A.A., M.E. I.K. and G.K. conceived and designed the experiments. I.F. and A.F. performed the Western blot assays. A.A. and M.E. performed the immunoprecipitation and LC-MS experiments, respectively. I.F., A.F., A.A., M.E. and G.K. analyzed and visualized the data. A.A., M.E. I.K. and G.K. wrote the relevant results sections of the paper. G.K. and I.K. obtained the funding for the work performed at the University of Sussex.

Experiments performed in Japan: D.H. conceived and designed the experiments. D.H. and Y.T. purified total RNA from *Lymnaea* tissues. D.H., R.M., and T.K. prepared the LymCBP recombinant protein and performed acetylation assay. H.S. and E.I. performed electrophysiological experiments. T.W. created the phylogenetic tree.

D.H. wrote the first version of the paper and all the authors contributed to the final version.

## Competing Interests

The authors declare no competing financial interests.

## Notes

### Competing Interest Statement

The authors have declared no competing interest.

### Summary of Updates

We have made a number of small amendments to the text.

